# Edison: measuring scaffolding accuracy with edit distance

**DOI:** 10.1101/2022.03.25.484952

**Authors:** Aakash Sur, William Stafford Noble, Shawn Sullivan, Peter Myler

**Affiliations:** Department of Biomedical Informatics and Medical Education, University of Washington; Department of Genome Sciences, University of Washington; Paul G. Allen School of Computer Science and Engineering, University of Washington; Phase Genomics; Seattle Children’s Research Institute; Department of Pediatrics, University of Washington

## Abstract

**Motivation:** The quality of reference genomes critically affects analyses of next generation sequencing experiments. During the construction of the reference genome, contigs are organized into their underlying chromosomes in the scaffolding step. Historically, the quality of scaffolding software has been difficult to evaluate in a systematic and quantitative fashion. To this end, we identified genomic edit distance as a compelling method for evaluating the quality of a scaffold.

**Results:** We present Edison, a Python implementation of the Double Cut and Join (DCJ) edit distance algorithm. Edison calculates the overall accuracy of a given scaffold using a reference genome and also provides scores for characterizing different aspects of the scaffolding accuracy, including grouping, ordering, and orientation. All metrics are calculated on a length-weighted basis, which rewards the correct placement of longer contigs over shorter ones. By creating 1000 random assemblies of the *S. cerevisiae* genome, we show that our scaffolding accuracy provides a more reliable metric than the commonly used metric, N50. Edison can be used to benchmark new scaffolding algorithms, providing insights into the strengths and weaknesses of each approach.

**Availability and Implementation:** Edison is available under an MIT license at https://github.com/Noble-Lab/edison.

## 1 Introduction

The reference genome of a species is the starting point for the many types of sequencing experiments. Accordingly, errors in the reference genome often propagate through subsequent analyses. The critical task of constructing the reference genome is handled by genome assemblers, which distill large sets of genomic reads into stretches of contiguous sequence. Ideally, a genome assembly contains chromosome-length sequences, but in practice only subregions of high confidence, known as “contigs,” can be independently assembled. Arranging these contigs in the correct chromosome grouping, order, and orientation is known as the *scaffolding problem*. Although the scaffolding problem remains challenging, advances in experimental methods such as chromatin conformation capture have allowed researchers to publish high quality scaffolds for historically difficult genomes Beier et al. [2017], Zhang et al. [2018, 2017].

Despite the importance of scaffolding to the assembly process, there is little agreement on how to assess the quality of a given scaffold. Since 1995, when the term “scaffolding” was introduced Roach et al. [1995], there has been a steady flow of new scaffolding algorithms, each with its own evaluation criteria (Supplementary Table 1). Typically, evaluation criteria fall into three categories: length metrics, visual plots, and error counts. The most common length metric is N50, which is the size at which contigs of equal or greater length cover half the assembly. Though useful, since this metric only characterizes the length of the scaffolds, it does not evaluate the placement of contigs, which can lead to overly aggressive scaffolders receiving higher scores for otherwise inaccurate scaffolds. Visual inspection using dot plots and linkage maps can confer a sense of agreement with a reference genome but does not yield quantitative measurements. In contrast, enumerating the errors in scaffolds creates quantifiable values, but there are many choices in defining these errors.

We suggest using the concept of edit distance, which measures the minimum number of edits (splitting scaffolds, joining scaffolds, moving contigs, and inverting contigs) required to fix misplaced contigs, to encompass all these flavors of accuracy. Edit distance has been studied in many contexts Berger et al. [2021], and in the field of evolutionary genomics it has generally been defined as the most parsimonious series of rearrangements in gene order that would explain the evolution of one species into another Hannenhalli and Pevzner [1995], Tesler [2002], Ozery-Flato and Shamir [2003], Yancopoulos et al. [2005]. In 2006, an elegant formulation called the Double Cut and Join (DCJ) model was introduced, which accounted for chromosomal fusions, fissions, translocations, and inversions Bergeron et al. [2006]. However, edit distance has rarely been applied to the evaluation of scaffolds Darling et al. [2010], Alonge et al. [2019], and to the best of our knowledge, there is no current software tool to perform this task.

We have developed Edison (<u>E</u>dit <u>Di</u>stance <u>S</u>caff<u>o</u>ldi<u>n</u>g), an open source Python that uses the DCJ edit distance between a scaffolded assembly and a high quality reference genome as a measure of scaffolding accuracy. This software package calculates the overall length-weighted edit distance (relative to the correct placement of contigs), along with individual scores for grouping, ordering, and orientation accuracy. By focusing strictly on contig placement instead of sequence error, we can disentangle errors in the genome assembly step and the scaffolding step. As such, our measurement of scaffolding accuracy allows researchers to benchmark existing scaffolders and test new algorithms against known genomes to gain a better understanding of performance compared to traditional metrics such as N50 and base level errors. Our findings show that on random permutations of the yeast genome, scaffolding accuracy better evaluates the state of an assembly compared to N50.

## 2 Results

Edison begins by breaking scaffolds into their constituent contigs at regions with a stretch of Ns. Next, contig sequences are aligned to the reference genome using MUMmer4 Marçais et al. [2018]. This alignment allows us to determine the optimal organization of contigs into scaffolds. To determine the edit distance, we compare this alignment-based scaffolding to the input scaffolds using the DCJ algorithm. The algorithm begins by constructing an adjacency graph that maps the positions of contigs in both assemblies onto a graph. The edit distance then becomes a simple relationship between the number of contigs and the number of even and odd paths in this graph (Supplementary Methods). The edit distance is converted into accuracy by taking the total length of correctly placed contigs compared to the overall length of all the contigs (Figure 1).

**Figure 1:**
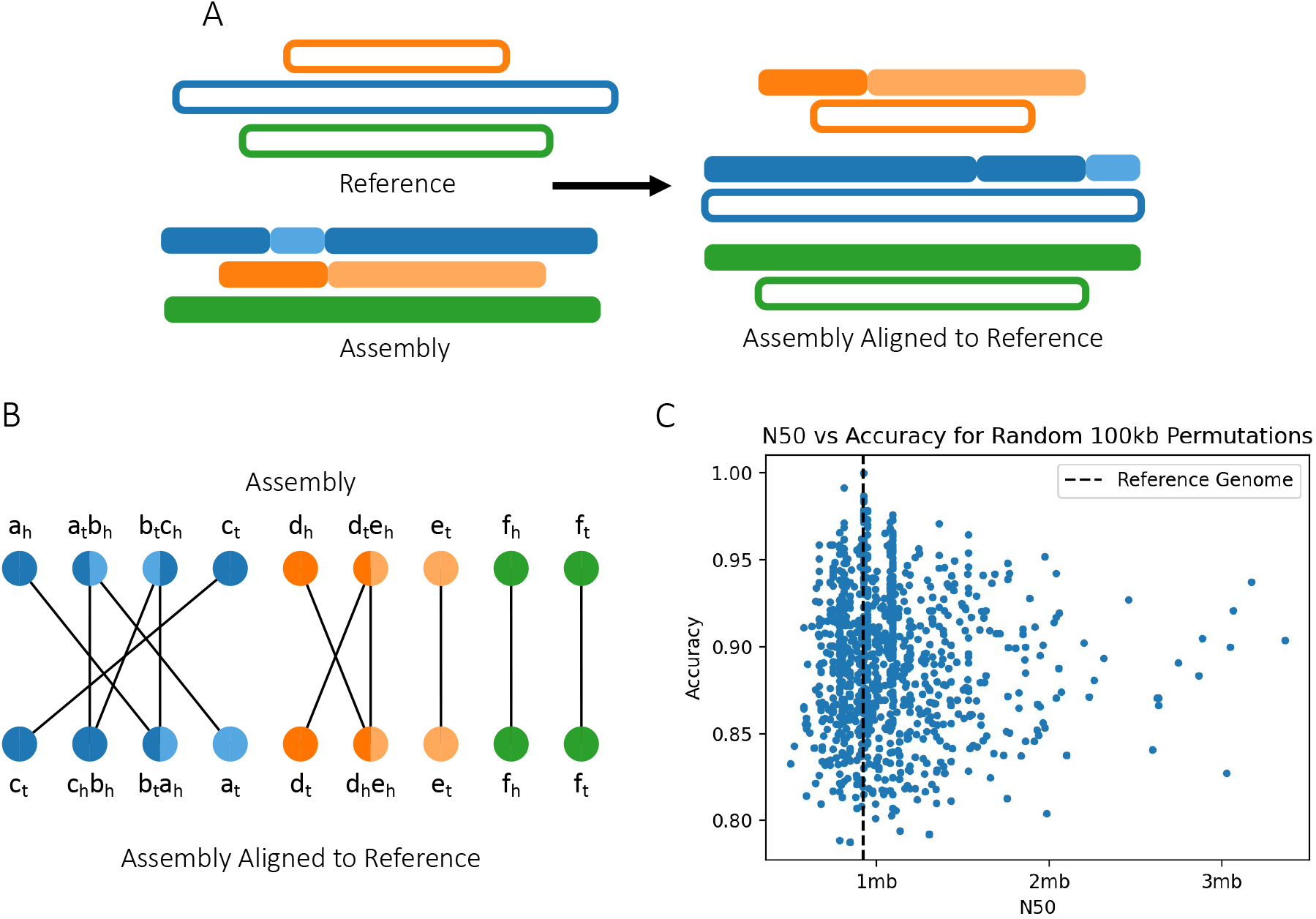
An overview of the Edison pipeline, and experimental results. A) Scaffolds in the assembly are first aligned to the reference genome to determine their optimal positions. B) We compare the original assembly and the assembly aligned to the reference by creating an adjacency graph. As described in the Double Cut and Join model, the adjacency graph can be used to compute the edit distance between these two layouts. C) Random permutations of the *S. cerevisiae* genome indicate that while the N50 can be artificially inflated, the accuracy cannot. The dashed vertical line represents the N50 of the reference genome.

To assist in the interpretation of the accuracy and edit distance, we break down three of its contributing factors, the grouping, ordering, and orientation scores. The grouping score represents what fraction of a scaffold belongs to a single chromosome, averaged across all scaffolds by length. The ordering score is the length weighted percent of contigs that are next to their expected neighbors. The orientation score is similar to the ordering score, but contigs are required to be in the correct orientation in addition to being ordered correctly. Finally, Edison also produces a visualization which displays the Mummer alignments of the contigs to further illuminate how a particular assembly compares to the reference genome.

In the presence of a reference genome, we observe that accuracy is a much better indicator of correctness than the N50 of the assembly. To demonstrate this, we simulated a genome assembly by splitting the *S. cerevisiae* reference genome into equal sized 100kb contigs and scaffolding them according to their true chromosomal assignments. We then randomly made between 0 and 30 permutations to the assembly by moving contigs, merging scaffolds, and breaking scaffolds to create 1,000 permuted assemblies. Several of these scaffolds had considerably higher N50s than even the reference genome, which would have made them attractive candidates in a *de novo* setting. However, the accuracy of these assemblies is demonstrably lower due to the incorrect joins required to create these longer scaffolds.

Developing new scaffolders is a challenging task which has to balance producing longer scaffolds with producing correctly joined scaffolds, and it has been shown that more aggressive scaffolding parameters accumulate more errors Coombe et al. [2018]. Because these scaffolders are most often used in the de novo setting, where a reference genome is absent, correctness is quite difficult to characterize. With Edison, we propose that researchers benchmark and test their scaffolders on assemblies of species with known reference genomes in order to compare performance between scaffolders and evaluate the strengths and weaknesses of competing algorithms.

