## Supplemental Text and Figures for "Edison: measuring scaffolding accuracy with edit distance"

March 18, 2022

Computing the accuracy of an assembly requires determining whether the grouping, order, and orientation of each contig within a scaffold matches its position in the corresponding chromosome of the reference genome. Here, we outline the steps necessary to do so, along with the equations to compute accuracy metrics.

#### 1 Mapping

All scaffolding methods output a FASTA file, in which contigs are joined in some particular order. We produce an "A Golden Path" (AGP) file to record this order, and then disassemble it into its constituent contigs by splitting scaffolds at runs of Ns (default: 10 Ns). NCBI [2019] Mummer 4 is then used to determine where contigs map on the reference genome. Alignments must have a minimum alignment length (default: 1000bp) and a minimum percent of the query sequence aligning to the reference (default: 20%), otherwise they are excluded from subsequent steps. From this coordinate map, we generate another AGP file representing the ideal scaffolding. Evaluation of scaffolding accuracy can then be computed solely from these two AGP files.

#### 2 Edit Distance

The method for computing the edit distance between two assemblies $A$ and $B$ begins by constructing the adjacency graph. In this bipartite graph, the two sets of vertices are the adjacencies in each genome, and edges connect vertices with overlapping adjacencies. The algorithm for creating the adjacency graph is outlined in greater detail in the original Double Cut and Join edit distance paper Bergeron et al. [2006]. The distance can then be computed as a function of $N$, the number of contigs, $C$, the number of cycles and $I$, the number of odd paths in the adjacency graph:

$$f_{\text{distance}} = N - \left(C + \frac{I}{2}\right) \tag{1}$$

In two assemblies that are identical, all contigs are involved in cycles of length two or odd paths of length two. This observation lets us compute a length weighted version of edit distance, our notion of accuracy, where the longest two contigs in each cycle and odd path are taken to represent the number of bases do not have to be moved:

$$f_{\text{accuracy}} = \text{len}(C) + \frac{\text{len}(I)}{2} \tag{2}$$

Intuitively, we can divide scaffolding into three tasks: grouping contigs into chromosomes, ordering contigs, and orienting them such that contiguous ends are touching. Though a scaffolder may not explicitly perform these tasks, they are always implicit in the output, allowing any scaffolder to be compared on these common fronts. Here, we further define the metrics for each of these sub-tasks.

#### 2.1 Grouping

To evaluate grouping performance, we must determine the degree to which scaffolds overlap with reference chromosomes. Suppose there exists some reference chromosome $A_i$ and some assembly scaffold $B_j$, then the intersection of these two sets of contigs are those contigs which belong to both the chromosome and the scaffold. We define the length weighted Jaccard index as the sum of contig lengths in the intersection divided by the sum of contig lengths in the union for any two sets:

$$J(A_i, B_j) = \frac{\text{len}(A_i \cap B_j)}{\text{len}(A_i \cup B_j)} \tag{3}$$

We then find the maximum length weighted Jaccard index for each reference chromosome by iterating through all the assembly scaffolds. These maximum Jaccard values are then weighted by the length of the reference chromosome it corresponds to, such that smaller chromosomes get weighed less. The sum of these weighted Jaccard maximums are then divided by the length of the reference genome:

$$f_{\text{grouping}}(A, B) = \frac{\sum_{i=0}^{|A|} \arg\max_j (J(A_i, B_j)) * \text{len}(A_i)}{\text{len}(A)} \tag{4}$$

Since the Jaccard index is a value between 0 and 1, the grouping score is also a value between 0 and 1.

#### 2.2 Ordering

The ordering performance can be evaluated by determining how many contigs were next to their expected adjacency. Given contigs $k$ and $l$ in the reference $A$, let their adjacency be $kl$. The two adjacencies $kl$ and $lk$ are then the same. The length weighted adjacency is then the length of $k$ plus the length of $l$. If we record all the length weighted adjacencies in the reference $A$ and the assembly $B$, then the ordering score is the sum of adjacencies in the intersection divided by the sum of adjacencies in the reference $A$:

$$f_{\text{ordering}}(A, B) = \frac{\text{len}((A_{edges}) \cap (B_{edges}))}{\text{len}(A_{edges})} \tag{5}$$

#### 2.3 Orientation

In a similar fashion, we can compute the orientation accuracy if we construct our adjacency set to also include orientation. Here, we retain which end of each contig is adjacent when recording the set of adjacencies such that contigs $i$ and $j$ might create an edge $i_h j_t$ indicating the head of $i$ is contiguous with the tail of $j$. Again, the adjacency $i_h j_t$ is equivalent to $j_t i_h$. Then the orientation score is the sum of adjacencies in the intersection divided by the sum of adjacencies in reference $A$:

$$f_{\text{ordering}}(A, B) = \frac{\text{len}((A_{edges'}) \cap (B_{edges'}))}{\text{len}(A_{edges'})} \tag{6}$$

### 3 Software

We implement the methods in a python package which is freely available under the MIT license (`https://github.com/Noble-Lab/2021_asur_scaffolding`). It takes in two inputs, a reference FASTA and scaffolded FASTA file, and chiefly produces five metrics of accuracy. While running, it also produces a plot of how the contigs map to the reference genome and the scaffolds to which they belong (Supplementary Figure 1). Additionally, it produces two AGP files representing how the assembly places contigs and how they ought to be placed to most closely match the reference.

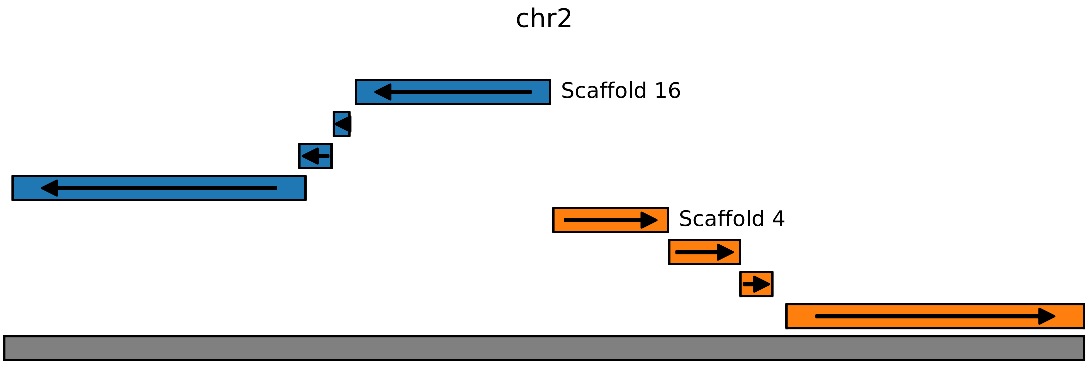


Figure 1: **Visualizing contig alignments to a reference genome.** Here, each row represents a new contig, where the horizontal coordinates indicate where on the reference chromosome they aligned, the arrow indicates the orientation of alignment, and the color corresponds to scaffold membership.

| | Year | N50 | Visual Plots | Misassembly | Grouping | Ordering | Orientation | Edit Distance |
|---|---|---|---|---|---|---|---|---|
| BambusPop et al. [2004] | 2004 | ✓ | | | | ✓* | ✓* | |
| SomaNagarajan et al. [2008] | 2008 | | | | | ✓* | | |
| AmosPhillippy et al. [2008] | 2008 | | | ✓ | | | | |
| MauveDarling et al. [2011] | 2011 | | ✓ | ✓ | | | | ✓* |
| SspaceBoetzer et al. [2011] | 2011 | ✓ | | | | | | |
| GageMagoc et al. [2013] | 2013 | ✓ | ✓ | ✓* | | | | |
| LachesisBurton et al. [2013] | 2013 | ✓ | ✓ | | ✓ | ✓ | ✓ | |
| Hunt'sHunt et al. [2014] | 2014 | | | | | ✓* | ✓* | |
| HirisePutnam et al. [2016] | 2016 | ✓ | | | | ✓* | ✓* | |
| 3d-dnaDudchenko et al. [2017] | 2017 | ✓ | ✓ | | | | | |
| ArksCoombe et al. [2018] | 2018 | ✓ | ✓ | ✓* | | | | |
| SalsaGhurye et al. [2019] | 2019 | ✓ | | | ✓* | ✓* | ✓* | |
| ScopLi et al. [2019] | 2019 | | | | | ✓* | ✓* | |
| LrscafQin et al. [2019] | 2019 | ✓ | | ✓* | | | | |
| SlrLuo et al. [2019] | 2019 | ✓ | | ✓* | | | | |
| AllhicZhang et al. [2019] | 2019 | ✓ | ✓ | | | | | |
| RagooAlonge et al. [2019] | 2019 | ✓ | ✓ | ✓* | ✓* | ✓* | ✓* | ✓* |
| LdscaffZhao et al. [2020] | 2020 | ✓ | ✓ | ✓ | | | | |
| Edison | 2022 | ✓ | | | ✓ | ✓ | ✓ | ✓ |

Table 1: **Scaffolding methods and the metrics they use to assess the accuracy.** "Visual plots" refers to any kind of visual analysis, such as dot plots, linkage plots, and circle plots. "Misassemblies" refers to the counting of structural variants such as inversions, deletions, substitutions, and translocations. *Indicates that the metric is a count and is not weighted by the length of the contigs.
